# Limits to the evolution of metabolic dependency in spatially structured microbial communities

**DOI:** 10.1101/2025.05.09.651753

**Authors:** Divvya Ramesh, Emanuele Fara, Franziska Oschmann, Giovanni Stefano Ugolini, Roman Stocker, Simon van Vliet, Martin Ackermann, Olga T. Schubert

## Abstract

In microbial communities, evolutionary processes can lead to loss of biosynthetic pathways, creating metabolic dependencies. The Black Queen Hypothesis suggests that such gene loss can confer a fitness advantage by reducing metabolic burden. However, how these dependencies evolve at the level of individual cells in spatially structured communities remains poorly understood. We used a combination of microfluidic single-cell imaging and mathematical modeling to examine the early fate of auxotrophic mutants within *E. coli* populations. We found that without external amino acids, auxotroph growth is strongly constrained by low amino acid leakage from wildtype neighbors, and further reduced when they form local clusters that drain this limited amino acid pool. A growth advantage was only observed when amino acids were added or when leakage from wildtypes exceeded a threshold. Together, our results reveal insights into determinants of mutant invasion fitness and the trade-offs between reducing metabolic costs and maintaining metabolic autonomy.

**Highlights:**

1. Single-cell imaging reveals how auxotrophs fare in spatially structured populations
2. Auxotrophs grow slowly due to low amino acid leakage from wildtype neighbors
3. Mutant clustering intensifies local amino acid depletion and reduces auxotroph growth
4. Model identifies leakage threshold required for auxotroph invasion from rare

## Introduction

Microbial life predominantly thrives in spatially structured habitats characterized by complex physical and chemical environments.^1–3^ In these settings, microbes form dense, interconnected metabolic networks, where they shape their microenvironment through the consumption and release of metabolites.^4,5^ Processes such as the leakage and uptake of costly metabolites, including amino acids, significantly influence growth, survival, and metabolic processes of both individual cells and the entire community.^6–9^ Most of the resulting interactions are mediated by diffusible molecules^1^ and their strength diminishes with increasing distance between cells.^10,11^ Consequently, spatial organization plays a critical role in defining the nature and extent of metabolic interactions within microbial communities.^12^

Due to the often large population sizes and short generation times, microbial populations are highly dynamic, continuously mutating and evolving. These evolutionary dynamics often lead to metabolic dependencies within communities. A common driver of such metabolic dependencies is the loss of biosynthetic pathways requiring cells to acquire the missing metabolites from the environment or other community members.^13–15^ This process is central to the Black Queen Hypothesis, which stipulates that such gene losses reduce metabolic burden and can thus confer a selective advantage when essential metabolites are available within the community.^16–19^ The loss of biosynthetic capabilities is indeed widespread across bacteria, suggesting that gene loss serves as a common adaptation to environments where the corresponding metabolites can be acquired from external sources.^20^ One possible source of such externally available metabolites is leakage by neighboring cells that continue to synthesize these compounds, including amino acids.^21^ Notably, many of these organic molecules cannot freely diffuse across the cell membrane. Instead, their release into the extracellular environment primarily relies on transport proteins, either passively via transporter back-reactions or actively to expel excess metabolites and maintain homeostasis.^21^ Experimental evolution studies have demonstrated that auxotrophic mutants lacking biosynthetic genes can gain a competitive advantage over prototrophic wildtypes when essential metabolites, such as amino acids, are externally supplied at sufficient concentrations.^20,22^ This advantage may arise because wildtype cells, despite regulatory mechanisms that adjust biosynthetic activity in response to external metabolite availability, may still incur residual metabolic costs due to incomplete repression of biosynthetic pathways.^23^ Gene loss could fully eliminate these costs, providing a direct fitness benefit to auxotrophic mutants.^20^

Despite growing evidence that auxotrophy is a common adaptation in microbial communities, how these dependencies evolve remains largely unknown.^24^ Amino acid auxotrophy, in particular, is among the most common forms of metabolic dependency observed in nature, possibly as a consequence of the high metabolic cost associated with amino acid biosynthesis.^24,25^ Previous studies have primarily examined auxotrophy in well-mixed conditions or inferred metabolic dependencies from comparative genomics, leaving open questions about their evolution in spatially structured communities.^17,20^ In such settings, metabolite exchange has been shown to be confined to immediate neighbors and strongly shaped by the spatial arrangement of cells.^26,27^ Studying how such dependencies evolve in spatially structured environments is essential for understanding the balance between reducing biosynthetic burden and relying on neighboring cells for essential nutrients.

Here, we investigated the first steps in the evolution of metabolic dependencies at single-cell resolution within spatially structured *Escherichia coli* populations, focusing on amino acid auxotrophy. To this end, we introduced individual cells with gene deletions in amino acid biosynthesis pathways into groups of wildtype cells. A microfluidic setup coupled with live-cell microscopy and automated image analysis allowed us to analyze single-cell growth rates and evaluate interactions between auxotrophic mutants and wildtype cells under different spatial configurations. Specifically, we addressed three key questions: (1) How do auxotrophic mutants grow compared to wildtype cells in the presence and absence of externally available amino acids? (2) How are auxotrophic mutants influenced by neighboring auxotrophs as they divide and form clusters? (3) How are wildtype cells influenced by neighbouring auxotrophs?

## Results

To understand how amino acid auxotrophs establish within wildtype populations, we studied the fate of single cells that lack genes for amino acid biosynthesis in spatially structured populations of prototrophic wildtype *E. coli*. Each experiment included a wildtype strain capable of synthesizing all amino acids and an auxotroph lacking the ability to synthesize a specific amino acid *(Figure. 1)*. The auxotrophs used in our study lacked the ability to synthesize methionine (ΔmetA), tryptophan (ΔtrpC), or proline (ΔproC), representing amino acids with different costs.^10,28^ To distinguish the strains in mixed populations, we chromosomally integrated a constitutively expressed gene for a red or green fluorescent protein into auxotrophs and wildtypes, respectively.

**Figure 1:**
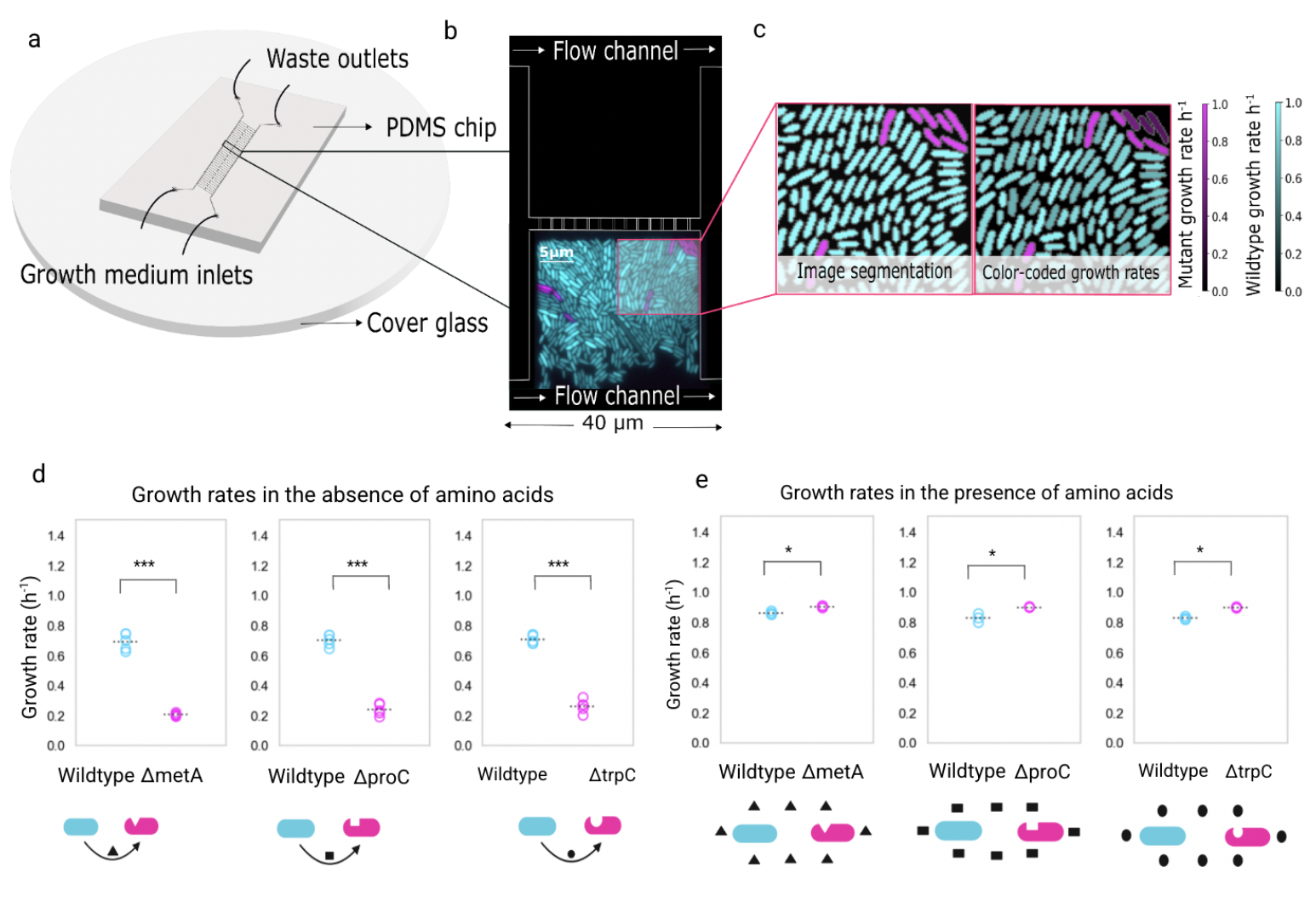
Experimental setup and growth rates of auxotroph and wildtype cells in the absence and presence of amino acids. (a) Schematic of the PDMS microfluidic chip bonded to a cover glass. Growth medium enters through inlet channels and flows through the central region of the device containing an array of growth chambers, with waste exiting via outlet channels. (b) Each chamber (40 × 40 × 0.8 µm) is open on one side to a main feeding channel (22 µm height, 100 µm width) and on the opposite side connected via narrow slits (0.5 µm wide, 3 µm long, spaced by 2 µm) to a secondary feeding channel. This design allows nutrients to enter the chambers from both sides, ensuring a more uniform supply across the chamber length. Cells were loaded only from the side open to the main feeding channel, while the opposite side received medium without cells. (c) Representative growth chamber showing wildtype cells (cyan) and auxotrophic mutants (magenta), false-colored based on fluorescent protein labeling. Single-cell growth rates were determined by segmentation and tracking of individual cells (left) over time, with growth rates visualized by color brightness (right, brighter colors indicating higher growth rates). (d) Mean growth rates of wildtype and auxotrophic cells for each of five biological replicates in the absence of externally provided amino acids. Data are represented as mean +/- SEM. In total, we analyzed the following numbers of cells: methionine: 30,879 wildtype cells and 337 auxotroph cells, proline: 30,444 wildtype cells and 295 auxotroph cells, and tryptophan: 36,748 wildtype cells and 346 auxotroph cells. Across all conditions, auxotrophic cells consistently exhibited significantly slower growth rates compared to wildtype cells, growing approximately three times slower (^***^ p < 0.001, paired t-test performed on the level of biological replicates). Schematic icons below each condition illustrate the interaction between wildtype and auxotrophic cells: curved arrows represent metabolic dependency, where auxotrophs rely on leaked amino acids from wildtype cells for growth. (e) Mean growth rates of wildtype and auxotrophic cells for each of five biological replicates in the presence of externally provided amino acids. Data are represented as mean +/- SEM. In total, we analyzed the following numbers of cells: methionine: 9,775 wildtype cells and 5,513 auxotroph cells, proline: 13,894 wildtype cells and 10,828 auxotroph cells, and tryptophan: 10,605 wildtype cells and 7,482 auxotroph cells. Across all conditions, auxotrophic cells showed consistently higher growth rates compared to wildtype cells (^*^ p < 0.05, paired t-test performed on the level of biological replicates).

We cultured these cells in a microfluidic setup that allowed us to monitor growth at the single-cell level through time-lapse fluorescence microscopy *(Figure. 1b)*. Using automated image analysis, we segmented and tracked single cells to measure their growth rates (*Figure. 1c* and *Supplementary Video 1-3*).

### Auxotrophic mutants grow three times slower than wildtype cells in the absence of amino acids

To model natural scenarios where spontaneous mutations generate auxotrophies, we seeded the microfluidic growth chambers with a small fraction of auxotrophic cells (∼10%) in a population of wildtype cells. These experiments were conducted in M9 minimal media supplemented with 0.1% glucose. Since no amino acids were supplied in the media, the auxotrophs’ growth relied on amino acids leaked by wildtype cells. Based on the Black Queen Hypothesis^19^, which posits that the loss of costly biosynthetic pathways reduces metabolic burden, one may hypothesize that auxotrophs achieve faster growth due to the uptake of amino acids released by the wildtype cells. However, contrary to this expectation, all three auxotrophs exhibited significantly slower growth across all conditions (*Figure. 1d*).

Wildtype cells grew at an average specific growth rate of 0.70 ± 0.001 h^−1^ (doubling time, t_d_ = 59.5 minutes; all growth rates are reported as mean ± standard error of the mean unless otherwise stated). Auxotrophs showed markedly reduced growth, with an average specific growth rate of 0.23 ± 0.009 h^−1^ (t_d_ = 181 minutes). Among the auxotrophs, the methionine auxotroph grew at 0.20 ± 0.008 h^−1^ (t_d_ = 204 minutes), the proline auxotroph at 0.24 ± 0.009 h^−1^ (t_d_ = 174 minutes), and the tryptophan auxotroph at 0.25 ± 0.009 h^−1^ (t_d_ = 166 minutes). This substantial difference in growth rates between wildtype and autotrophs was consistent across five independent replicate experiments (*Figure. S1*) and statistically significant (paired *t*-tests: methionine, *t* = 19.47, *p* < 0.001; proline, *t* = 123.07, *p* < 0.001; tryptophan, *t* = 18.74, *p* < 0.001). Pairwise comparisons between auxotroph strains showed no significant differences (*p* > 0.2 for all pairs), indicating that all three auxotrophs faced comparable growth limitations despite their reliance on different amino acids. These results suggest that amino acid leakage from wildtype cells is low and insufficient to support robust auxotroph growth in these conditions.

In contrast, when the corresponding amino acid was supplemented at a concentration of 150 µM, auxotrophs exhibited significantly higher growth rates than wildtype cells (*Figure. 1e*). Across all three supplemented conditions, auxotrophs reached an average specific growth rate of 0.90 ± 0.003 h^−1^ (t_d_ = 46.2 minutes), surpassing the wildtype average of 0.84 ± 0.003 h^−1^ (t_d_ = 49.8 minutes). Among the auxotrophs, the methionine auxotroph grew at 0.90 ± 0.003 h^−1^ (t_d_ = 46.1 minutes), the proline auxotroph at 0.90 ± 0.002 h^−1^ (t_d_ = 46.3 minutes), and the tryptophan auxotroph at 0.90 ± 0.003 h^−1^ (t_d_ = 46.3 minutes). Paired *t*-tests confirmed significant differences between wildtype and auxotrophic strains (*Figure. S1*): methionine (*t* = -4.32, *p* = 0.050), proline (*t* = -4.55, *p* = 0.045), and tryptophan (*t* = -16.02, *p* = 0.004). As before, pairwise comparisons among auxotrophs revealed no significant differences (*p* > 0.2 for all pairs), indicating that all three auxotrophs benefited similarly from external amino acid supplementation. The enhanced growth of auxotrophic mutants in supplemented environments likely results from the loss of biosynthetic pathways genes, which can reduce metabolic burden.^17,29^

Overall, these findings highlight the pivotal role of the availability of amino acids in determining the evolutionary dynamics of auxotrophy. In the absence of external sources of amino acids, auxotrophic mutants cannot invade *(Figure. S7)* because they grow slower than the surrounding wildtype cells, presumably because amino acid leakage by wildtype cells is low. However, when amino acids are present from external sources, auxotrophs benefit from their intrinsic growth benefit and can invade from rare.

### Mutant clustering negatively affects growth of mutants and neighbouring wildtype cells

The evolutionary dynamics of spatially structured populations depend not only on the growth of individual mutant cells but also on how these mutants grow once they expand into small clusters. For auxotrophic mutants, which rely on externally available amino acids, clustering may further deplete local resources, as all cells in the cluster consume amino acids but do not produce them. We hypothesized that clustering intensifies competition for these limited metabolites, ultimately restricting the growth of auxotrophs within the cluster.

To test this, we defined a circular neighborhood with a 5-micrometer radius around each cell, encompassing its nearest neighbors. We then calculated the fraction of auxotrophic mutants within this neighborhood (*Figure. 2a*).^10^ This radius was chosen as it reflects the approximate length scale over which amino acids were previously found to be exchanged^10^, and was also predictive for mutant growth in our own experiments.

**Figure 2:**
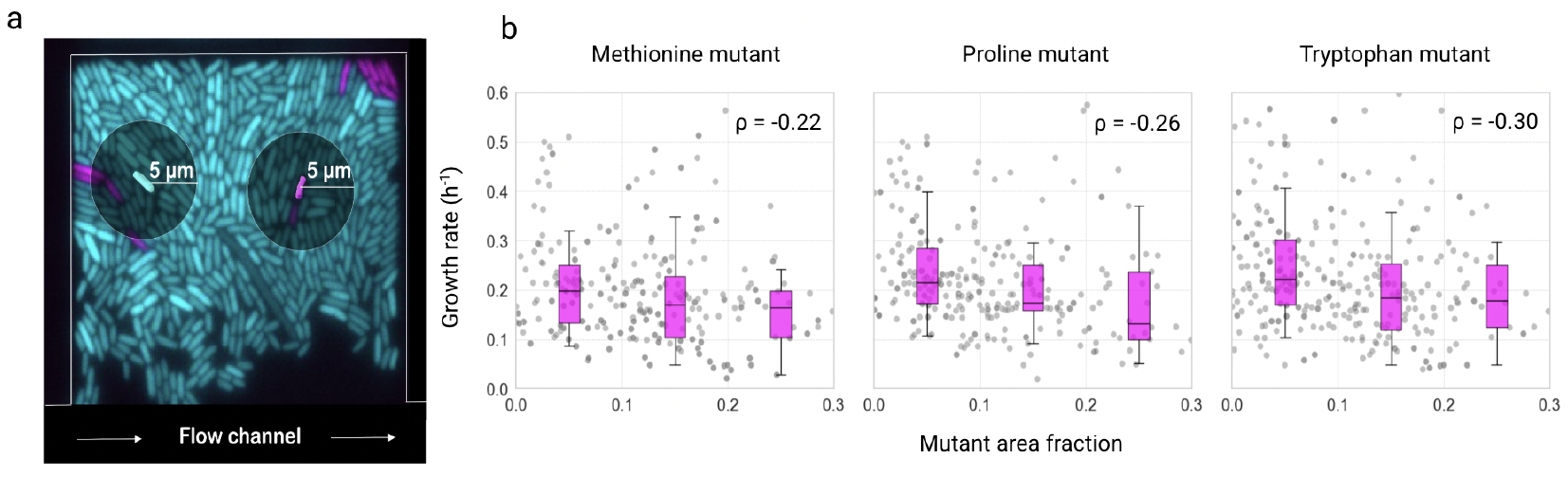
Local clustering of mutant cells influences the growth of mutants in the absence of externally supplied amino acids. (a) Representative fluorescence image showing auxotroph cells (magenta) and wildtype cells (cyan), with overlaid circular regions of interest (5 µm radius) centered on a focal auxotroph and wildtype cell. These regions define the local neighborhood used to compute the surrounding fraction of auxotrophs for each focal cell. (b) Relationship between the local neighborhood composition and auxotroph growth. Each data point represents a single cell. Box plots show the distribution of growth rates across fixed bins of the local auxotroph fraction. While the data was binned for visualization purposes, all statistical analyses were performed on the original, continuous data. Auxotrophs surrounded by a higher fraction of other auxotrophs exhibited significantly reduced growth, consistent with stronger negative selection at higher local auxotroph frequencies in spatially structured populations.

Our analysis showed that clustering indeed reduced the growth of auxotrophs, with auxotrophs growing more slowly when surrounded by other auxotrophs. This effect was quantified by calculating Spearman’s rank correlation coefficients between the growth rate of individual auxotroph cells and the fraction of their neighbors in the local neighborhood that were also auxotrophs. Combined across replicate experiments, all auxotrophs showed significantly negative correlations (methionine auxotroph ρ = -0.22, Proline auxotroph ρ = -0.26, Tryptophan auxotroph = ρ = -0.30; permutation test: *p* < 0.001) (*Figure. S3*). For better visualization, we binned the data into quartiles based on local mutant fraction (*Figure. 2b*). This revealed a clear trend where a 10% increase in the fraction of auxotrophic mutants within the local neighborhood was associated with an average decrease in growth rates of 7% for methionine auxotrophs, 12% for proline auxotrophs, and 9% for tryptophan auxotrophs.

These results indicate that even small increases in local auxotrophic mutant density can reduce their growth further due to competition for the low amounts of amino acids leaked from surrounding wildtype cells. This exemplifies negative-frequency dependent selection in its broader sense, where mutant fitness declines as their frequency increases, albeit without a sign-change, as auxotrophs are disadvantaged at all frequencies compared to wildtype.^29,30^ With spatial structure, this effect is compounded: auxotrophs form localized clusters, which leads to sharper reductions in amino acid availability within their immediate microenvironments. As a result, spatial organization amplifies negative selection by creating microenvironments where auxotrophs face stronger local competition for a depleted resource, and this disadvantage intensifies as their local frequency increases. In spatially structured populations, what matters is the local - not global - frequency of cell types. Due to short-range interactions and kin clustering, local frequencies can shift rapidly over just a few cell divisions, even while global frequencies remain largely unchanged. Thus, spatial structure accelerates the feedback loop between auxotroph clustering and resource depletion, reinforcing their growth disadvantage.

Given the significant effect of neighborhood composition on auxotroph growth, we next asked whether wildtype cells are also negatively affected by nearby auxotrophs. If auxotrophs impose a metabolic burden on wildtype cells by depleting the shared amino acid pool, this could reduce wildtype growth. Wildtype cells can normally reabsorb some of the amino acids they leak, but when auxotrophs are nearby acting as strong sinks, this recycled uptake is disrupted, potentially forcing wildtype cells to synthesize more amino acids to meet their own needs. To test this, we conducted a similar neighborhood analysis for focal wildtype cells.

Similar to auxotrophs, wildtype cells also exhibited reduced growth when surrounded by higher local densities of auxotrophs. While the effect was milder compared to that observed in auxotrophs, it was consistently significant across amino acids (*Figure. S2*). Combined across replicates, we found a modest but significant negative correlation between wildtype growth and local auxotroph density (methionine: ρ = –0.12; proline: ρ = –0.11; tryptophan: ρ = –0.09; *p* < 0.05 for all; *Figure. S3*). These findings indicate that auxotrophs not only experience fitness costs from clustering but also impose measurable burdens on neighboring wildtype cells, likely through localized depletion of shared amino acids.

As expected, when external amino acids were added, the effect of local mutant density on growth rate disappeared because the dependency on local amino acid exchange and any associated spatial effects on growth were eliminated (*Figure. S5*).

### Low amino acid leakage limits the invasion of auxotrophic mutants despite potential growth benefit

Our experimental results show that if amino acids are not externally provided, auxotrophic mutants cannot invade populations of wildtype cells *(Figure. S7)*. A plausible intuitive explanation for this outcome is that the intrinsic growth benefit of the mutants is not high enough to compensate for the low rates at which they receive amino acids through leakage from the wildtype. To probe this intuition, and better understand the conditions under which auxotrophy could evolve, we used a mathematical model developed by van Vliet and colleagues (Figure. *3a*).^31^ The model incorporates amino acid leakage and uptake rates as well as the growth advantage provided by auxotrophy, enabling us to analyse how these parameters influence the growth of individual auxotroph cells in a wildtype population (see Materials and methods). To parameterize the model, we combined two experimental datasets: first, the relative growth benefit of auxotrophs when amino acids are abundant (derived from supplementation assays) and, second, the maximum auxotroph growth rate observed when surrounded entirely by wildtype neighbors (*Table S1*). Using the latter, we calibrated the model by identifying the amino acid leakage rate that yields a predicted auxotroph growth rate matching our experimental observations. This approach allowed us to estimate leakage fluxes for proline, tryptophan, and methionine.

The model reveals how the growth rate of auxotrophs, relative to wildtype cells, changes with increasing amino acid leakage rates (*Figure. 3b*) under conditions where wildtype cells are the sole source of amino acids. As expected, higher leakage rates increase the relative growth rate of auxotrophs. The contour line within the heatmap delineates the conditions under which auxotroph and wildtype cells grow at equivalent rates, highlighting the critical leakage threshold above which auxotrophs have an advantage over wildtypes in the absence of externally supplied amino acids. These findings underscore that while auxotrophy can theoretically confer a growth advantage by reducing metabolic costs, this advantage is diminished under conditions of low amino acid leakage as we observe for the three amino acids studied here (*Figure 3b*). This highlights the delicate balance between reducing metabolic burden and becoming dependent on limiting metabolites.

**Figure 3:**
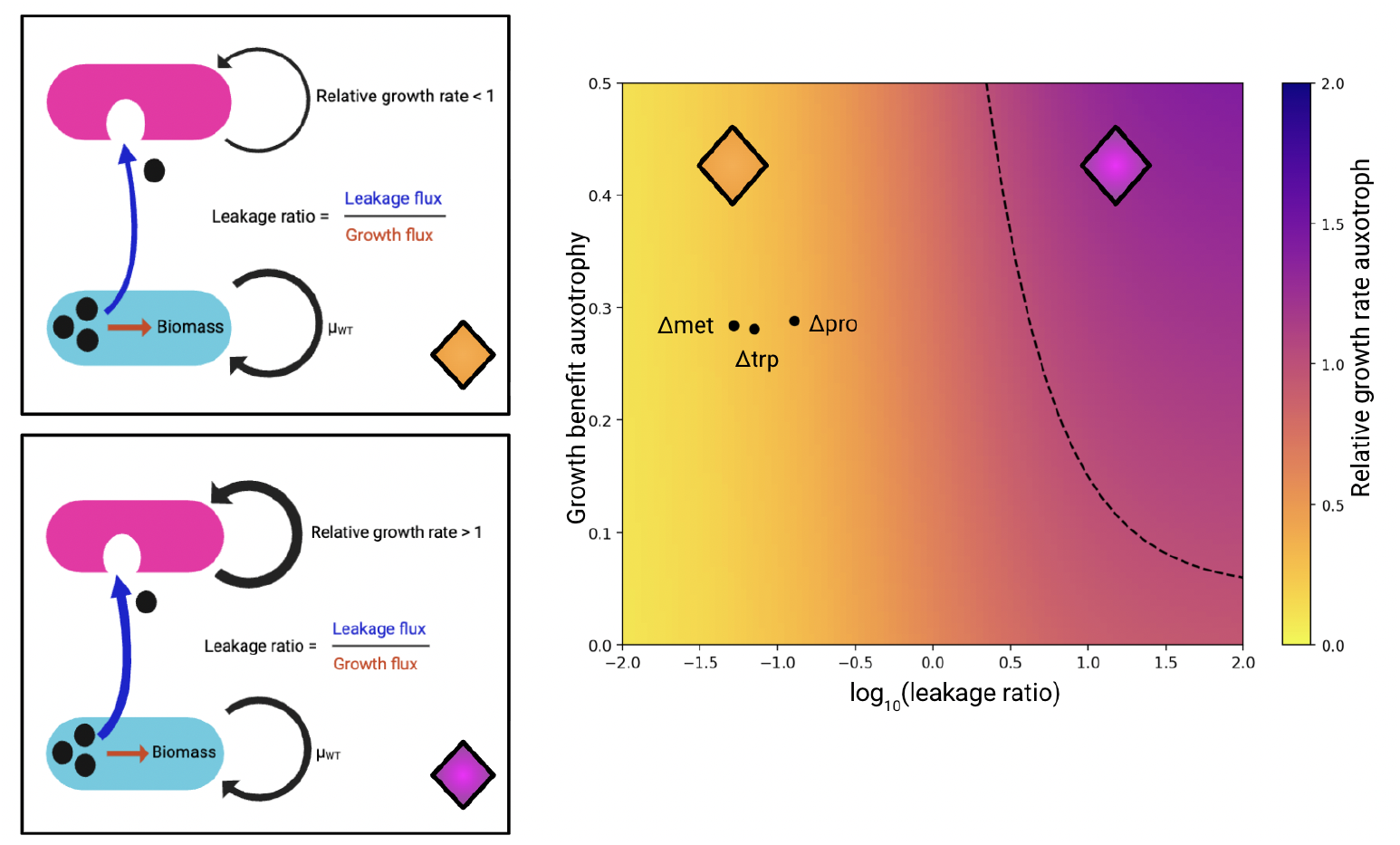
A mathematical model reveals tradeoffs for auxotroph evolutionary dynamics. (a) Schematic representation of the model capturing amino acid leakage by wild-type cells and uptake by auxotrophic mutants. Wild-type cells (cyan) synthesize an internal pool of amino acids: most is retained for biomass production (growth flux), while a small fraction is lost through leakage (leakage flux). Auxotrophic mutants (magenta) cannot synthesize the amino acid and must import the leaked pool to grow. The leakage ratio-defined as leakage flux divided by growth flux—quantifies the proportion of resources available to the mutant. Two scenarios are shown, corresponding to the parameter combination marked with an orange and a cyan diamond in panel (b) In the low-leakage scenario (top), the leakage ratio is small, so the mutant’s relative growth rate remains below that of the wildtype. In the high-leakage scenario (bottom), an increased leakage ratio enables the mutant to outgrow the wildtype. (b) The heatmap illustrates how metabolite leakage rate from wildtype cells^32^ and the associated growth benefit of auxotrophy (the growth benefit is defined as 1 + s, where s is the proportional increase in the auxotroph’s growth rate relative to the wildtype) influence the relative specific growth rate of amino acid auxotrophs. The dashed contour line marks the conditions under which auxotrophs and wildtype cells achieve equivalent growth rates. Highlighted data points represent experimentally derived leakage rates^32^ and growth benefits for methionine (Δmet), proline (Δpro), and tryptophan (Δtrp) auxotrophic mutants.

## Discussion

The aim of this study was to provide insights into how newly emerging mutants with metabolic dependencies evolve in a spatially structured population of cells, using amino acid auxotrophs as a model system. Our findings demonstrate that amino acid auxotrophs grow significantly slower than wildtype cells in spatially structured environments without external amino acids. While the Black Queen Hypothesis suggests that losing metabolic pathways can reduce metabolic costs and confer a growth advantage,^19^ our results highlight that realising this advantage strongly depends on environmental conditions. Specifically, low amino acid leakage from wildtype cells severely restricts auxotroph growth, creating a double disadvantage: first, auxotrophs inherently grow more slowly due to limited metabolite access; second, their growth is further suppressed when cells cluster, as competition for amino acids intensifies. However, when amino acids are supplied externally at high concentrations, auxotrophs can outgrow wildtype cells, underscoring the crucial role of environmental nutrient availability.

In spatially structured systems, the fate of a cell type depends not only on its growth but also on its initial location and physical properties like adhesiveness and motility. In our study, the only difference between WT and mutant cells is in their ability to synthesize a single amino acid, which is why we do not expect any systematic difference between these latter aspects, and we thus assume that any differences in fitness, averaged over many microfluidic growth chambers, are solely due to differences in growth rate. Here, we interpret single-cell growth rates as a proxy for short-term ecological performance, which allows us to identify the immediate constraints faced by newly arisen mutants.

Together, our experiments and modelling suggest that wildtype amino acid leakage rates would need to be much higher, by a factor of approximately 25, to enable auxotrophs to outcompete wildtypes based solely on their reduced biosynthetic burden. This positions all three auxotrophs below the critical threshold (*Figure 3b*), where the benefit from taking up leaked amino acids offsets their biosynthetic cost, explaining their inability to invade wildtype populations in the absence of externally supplied amino acids. This underscores a key limitation of the Black Queen Hypothesis: while it explains the evolution of metabolic dependency in environments with high metabolite availability or exchange, auxotrophy can be detrimental in environments with limited resource leakage.^33^ In such settings, auxotrophs’ reliance on external amino acids constrains their growth and evolutionary success.

Interestingly, one might expect auxotrophies of biosynthetically costly amino acids to differ in their fitness consequences compared to those of less costly amino acids. Yet we observed very similar growth rates across methionine, proline, and tryptophan auxotrophs, both in the absence (*Figure. 1d*) and in the presence *(Figure. 1e*) of externally supplied amino acids. In the absence of supplementation, the growth rates of auxotrophs in our experimental system are probably determined less by the intrinsic energetic cost of the missing pathway and more by the uniformly low leakage of amino acids from wildtype cells, which limits external supply and constrains all three auxotrophs similarly. When amino acids are supplied at saturating concentrations, biosynthetic pathways are downregulated and the growth benefit of auxotrophs likely results from eliminating residual expression.^34,35^ Our data suggests that the costs of residual expression are similar across the three amino acids that we tested. We see two factors that may contribute to this effect. First, biosynthetic pathways with higher expression costs are regulated more tightly than pathways with lower expression costs,^36^ leading to similar metabolic costs of residual expression across different amino acid biosynthetic pathways and hence a similar benefit upon their loss in the auxotrophs. Second, differences in per-molecule biosynthetic costs of amino acids are compensated by differences in amino acid flux, with more costly amino acids being produced in smaller quantities,^8^ resulting in similar overall costs across amino acids and hence a similar benefit for the auxotrophs.

Our findings highlight a key feature of structured environments in shaping auxotroph fitness. Whereas in well-mixed systems, such as planktonic microbial communities, auxotrophs can access nutrients uniformly and their growth is primarily limited by the overall availability of amino acids^37^, spatially structured environments, such as biofilms, impose physical constraints on metabolite exchange, intensifying local nutrient competition and amplifying the negative effects of mutant clustering.^11,38,39^ This may help explain why auxotrophy is common in environments with reliable metabolite exchange but is rare in environments where resources are scarce or inconsistently available.^40,41^ In other words, auxotrophy is difficult to sustain when cells must rely solely on passive metabolite leakage from neighbors.

However, auxotrophy may be more prevalent in environments where additional sources of amino acids are available. One important example is cell lysis, which releases intracellular contents, including amino acids, into the environment. Lysis can be induced by bacteriophages^42^, contact-dependent killing mechanisms like the type VI secretion system^43^, or other forms of cell death. In environments with frequent lysis, transient nutrient surges could be exploited by auxotrophs. For instance, phage-mediated lysis in marine ecosystems is estimated to kill 20-40% of all marine bacteria daily, releasing amino acid rich-dissolved organic matter^44^ that supports auxotrophic populations.^41^ This highlights how external factors, such as phage activity or other lysis-inducing mechanisms, can reshape the ecological dynamics of auxotrophs within microbial communities. In addition to such external sources, intracellular metabolic imbalance could also lead to the externalization of amino acids.^21^ Under balanced growth, leakage and growth fluxes typically scale proportionally, such that slower growth alone does not increase leakage. However, when metabolism becomes unbalanced, for example, under nutrient limitation or stress, cells may accumulate intermediates or experience overflow metabolism, resulting in active excretion of metabolites and a higher effective leakage ratio.^21,46^ Such stress-induced excretion could therefore represent another route by which amino acids become transiently available.

Beyond the specifics of our experimental system, our findings provide broader insights into the evolutionary and ecological dynamics of auxotrophs in microbial communities. While auxotrophs face intrinsic growth limitations in spatially structured environments due to low metabolite leakage by community members,^10^ external nutrient inputs may help alleviate these constraints, potentially supporting their persistence in natural settings. For instance, in the gut microbiome, where host-derived nutrients are abundant, auxotrophies are highly prevalent,^25^ presumably because auxotrophs can bypass the need for cross-feeding. Similarly, auxotrophs thrive in soil and aquatic ecosystems where the breakdown of organic matter provides a consistent supply of amino acids.^24^ Collectively, these observations emphasize that the widespread occurrence of auxotrophies is not solely driven by metabolic trade-offs but is strongly influenced by the ecological context and the availability of external amino acids.

An important consideration that remains unexplored in our study is the role of interspecies interactions in shaping auxotroph fitness. While our work focused on interactions within a single species (*E. coli*), microbial communities in nature consist of diverse species with complex metabolic interdependencies.^47^ In such communities, cross-species interactions could alter the dynamics of amino acid exchange and auxotroph fitness in several ways. For example, different species may exhibit varying levels of amino acid leakage or uptake efficiency, potentially creating new niches for auxotrophs.^16,41,48^ In particular, the presence of a shared mutualist can stabilize the coexistence of multiple auxotrophs by limiting the extent of resource monopolization.^49^ Moreover, amino acid release is not always passive; many species actively externalize metabolites to maintain intracellular metabolic balance. This form of active externalization could be especially relevant in multispecies communities, where species-specific metabolic strategies result in distinct externalization profiles that shape nutrient availability for surrounding cells.^21^

The widespread occurrence of amino acid auxotrophies in nature can be reconciled with our results by considering two distinct evolutionary processes. First, adaptive evolution may favor gene loss when environmental conditions reliably supply amino acids from external sources. In such settings, eliminating biosynthetic pathways can reduce proteome investment, freeing up cellular resources like ribosomes and lowering energy and redox demands, thereby enhancing growth.^25,41^ However, our findings show that in environments where amino acid availability is limited and leakage rates are low, this adaptive advantage cannot be realized, and auxotrophs are at a growth disadvantage. Second, mutation accumulation may contribute to the prevalence of auxotrophy in nutrient-rich environments.^50^ When external amino acids are consistently available, mutations disrupting biosynthetic pathways are not strongly selected against, allowing conditionally deleterious mutations to accumulate over time.^50^ In such cases, auxotrophs persist not because of an immediate growth advantage, but because the fitness cost of losing biosynthetic capacity is masked by the environment. Together, these processes - adaptive gene loss under abundant nutrient conditions and neutral mutation accumulation - highlight how ecological context fundamentally shapes the evolution and maintenance of auxotrophies, helping to explain their high prevalence despite the growth limitations we observe under nutrient-poor conditions.

In summary, our study shows that, depending on environmental amino acid availability, low amino acid leakage can be a significant barrier to the evolution of auxotrophy, even when gene loss offers a potential metabolic advantage. The success of auxotrophs depends strongly on environmental context, and in particular on access to essential metabolites from external sources. Our work provides a foundation for understanding how spatial structure and nutrient availability shape auxotroph fitness in single-species systems. Extending this work to multi-species communities will offer deeper insights into the ecological and evolutionary dynamics of auxotrophies in natural environments, where cross-species interactions are prevalent.

## STAR methods

### EXPERIMENTAL MODEL

#### Bacterial strains

Experiments were performed using *E. coli* MG1655-derived strains with the following fluorescent labeling scheme: All auxotrophic mutants were genomically labeled with mCherry, and the wildtype strain with sfGFP. The three auxotrophic strains ΔtrpC-mCherry (MG1655 *trpC*::frt, PR-*mCherry*), ΔproC-mCherry (MG1655 *proC*::frt, PR-*mCherry*), Δ*metA*-*mCherry* (MG1655 *metA*::frt, PR-*GFP*) were constructed by introducing the respective kanamycin resistance cassettes from the Keio collection^51^ into TB205^52^, a strain constitutively expressing mCherry from the chromosomal lambda promoter (PR), using lambda red-mediated recombination.^53^ The wildtype strain WT-GFP (MG1655, PR-*GFP*) was derived from TB204, which expresses sfGFP from the same promoter. For each mutant, the kanamycin cassette along with flanking homology arms was PCR-amplified and introduced into TB205 by electroporation. Successful transformants were confirmed to be auxotrophic based on their inability to grow in minimal media lacking the corresponding amino acid, both in liquid culture and on agar plates. Kanamycin cassettes were subsequently excised using FLP recombinase expressed from plasmid pCP20.^53^ The PCR primers used are tabulated below.

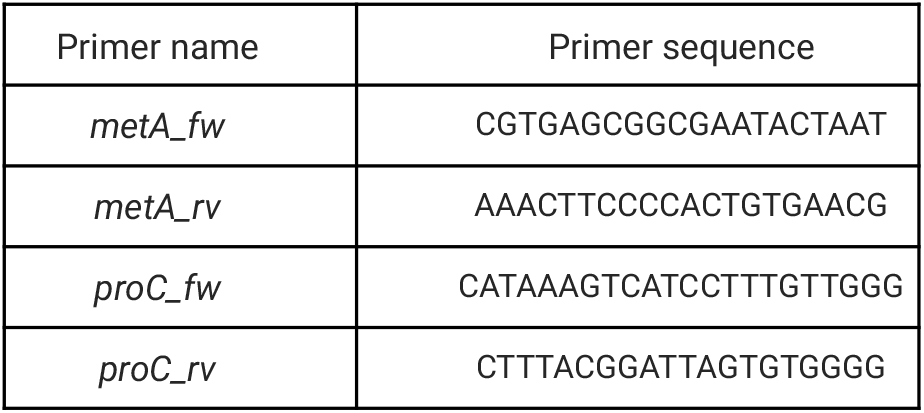
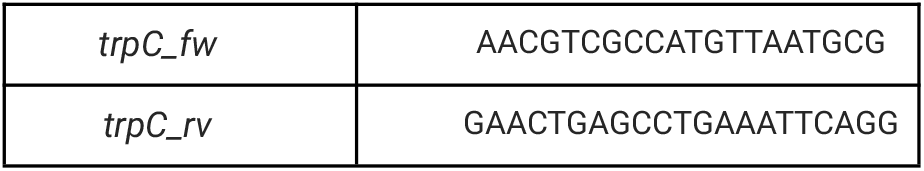

To confirm that the expression of different fluorescent markers does not affect growth, we compared the two wildtype reference strains TB204 (expressing sfGFP) and TB205 (expressing mCherry) in monoculture. Growth curves obtained in minimal medium with and without amino acid supplementation showed nearly identical growth dynamics and maximal growth rates for both strains (*Figure. S8*). This confirms that the fluorescent marker genes used throughout this study do not differentially affect growth in the wildtype background. Consistently, auxotrophic derivatives of these strains also exhibited comparable growth curves, indicating that the observed phenotypic differences in mixed communities arise from the introduced auxotrophies rather than from the fluorescence markers.

## METHOD DETAILS

### Media and growth conditions

Three sets of experiments were conducted, each pairing the wildtype strain with a specific amino acid auxotroph. Monocultures were started from single colonies grown on rich lysogeny broth (LB) agar plates and incubated overnight at 37°C in a shaker at 220 rpm. The growth medium was M9 minimal medium (47.76 mM Na_2_HPO_4_, 22.04 mM KH_2_PO_4_, 8.56 mM NaCl, 18.69 mM NH_4_Cl), supplemented with 1 mM MgSO_4_, 0.1 mM CaCl_2_, and 0.1% glucose. For auxotrophs, the medium was additionally supplemented with 150 µM of the required amino acid. To confirm that auxotrophs could not grow in the absence of externally supplied amino acids, they were cultured in M9 minimal medium with 0.1% glucose without amino acid supplementation, where no growth was observed.

### Microfluidic device fabrication

The microfluidic device consisted of an array of adjacent chambers (40 µm × 40 µm × 0.80 μm) flanked on opposite sides by feeding channels measuring 22 μm in height and 100 μm in width (*Figure. 1a*). The chambers are connected by small gaps of 0.5 μm in width, 3 μm in length and spaced by 2 μm. Nutrients enter each chamber from both sides, enabling a more uniform supply across the chamber length. Cells were loaded only from one side, leaving the opposite chamber empty but filled with media. The master mold was fabricated by direct-laser writing into SU-8—a negative photoresist that crosslinks where exposed to light—using a Heidelberg DWL66+ on two sequential spin-coated layers (SU8-2001 and SU8-2025) (Pacheco, Ugolini et al., submitted).

The microfluidic chips for the experiments were prepared by replica molding of Polydimethylsiloxane (PDMS, Sylgard 184, Dow Corning), mixed at a 1:10 ratio, onto the master mold. The PDMS was cured at 80ºC for 45 minutes, cut into chips of 3.5 cm × 3.5 cm, with 0.5 mm holes punched on both ends for loading cells and medium provision. For assembly, the PDMS chips and 50 mm diameter glass coverslips (Menzel-Gläser) were plasma-treated for 1 minute, brought into contact, and then placed on a 100°C hot plate for bonding. All chips used in this study were bonded on the day before performing the experiment.

### Single-cell growth experiments using time-lapse microscopy

Single-cell growth experiments were conducted using a microfluidic device and time-lapse microscopy to track bacterial growth under controlled conditions. Monocultures of wildtype and auxotrophic strains were grown to mid-exponential phase, concentrated approximately 100-fold by centrifugation, and washed four times in M9 minimal medium to remove residual amino acids. Auxotroph and wildtype cells were mixed at a 1:9 ratio to simulate the emergence of a few auxotrophs in a wildtype population. The mixture was loaded into microfluidic growth chambers via pipetting, ensuring an initial composition of ∼90% wildtype and ∼10% auxotroph cells.

After loading the chambers with cells, M9 minimal medium without amino acids was supplied continuously using a syringe pump (NE-300, NewEra Pump Systems) connected to a 50 mL syringe via Microbore tubing (ID 0.76 mm, OD 2.29 mm, Saint-Gobain Tygon™). The medium was delivered at a flow rate of 0.4 mL/hour to prevent bacterial accumulation in the main channel. Polytetrafluoroethylene tubing (ID 0.3 mm, OD 0.8 mm, Adtech) connected the syringe setup to the polydimethylsiloxane (PDMS) microfluidic chip inlet.

Time-lapse imaging was performed using Olympus IX81 inverted microscopes equipped with a 100× NA1.3 oil objective and an ORCA-flash 4.0 v2 sCMOS camera. Fluorescent imaging was achieved using an X-Cite120 lamp and Chroma 49000 series filters for GFP and mCherry. Focus was maintained by the Olympus Z-drift compensation system, and the setup was controlled using Olympus cellSens software. Imaging was conducted at 37°C, with multiple positions on the same microfluidic device captured every five minutes in GFP, mCherry, and phase contrast channels. The experiment was run for ∼12 hours. Images were initially acquired in ets format and later converted to tiff for analysis.

For each amino acid auxotroph, five independent experiments (“biological replicates”) were conducted using different microfluidic chips and different batches of media. Each biological replicate corresponds to one channel in a microfluidic chip, with an average of five chambers analyzed per channel. The final number of cells in each dataset is provided in Table S2.

## QUANTIFICATION AND STATISTICAL ANALYSIS

### Automated image analysis

Image analysis was conducted using a custom-made software, MIDAP (https://github.com/Microbial-Systems-Ecology/midap, version 0.3.18). The tiff images were processed in MIDAP, where an interactive cutout tool was used to select from each tiff stack the region of interest, excluding approximately 8 µm closest to the main channel. Then, cells were segmented in the GFP and mCherry channels using a high-precision morphology algorithm called Omnipose^54^, followed by cell tracking with STrack^55^, which calculates pixel overlap between consecutive time points to track individual cells. MIDAP’s outputs included processed images, labeled and binary segmentations, cell counts per frame, and a comprehensive tracking file. The tracking file contained detailed data such as cell IDs, positions, areas, axis lengths, and fluorescence intensities for each channel, facilitating in-depth analysis of cellular growth and fluorescence dynamics over time.

### Estimation of single-cell growth rates

Single-cell growth rates for wildtype and auxotrophic mutants were calculated using cell length (major axis length) as a proxy for cell size. Cell dimensions, measured in pixels, were converted to micrometers (µm) using a pixel size of 0.065 µm. Growth rates were estimated assuming exponential growth, with cell lengths log-transformed and linear regression applied to the log-transformed (ln) values over the full lifetime of each cell. The resulting values were then converted to specific growth rates per unit of biomass (h^−1^). To ensure accuracy, filtering criteria were applied. Cells were excluded if their relative length change exceeded ±15% between consecutive frames, only tracks with at least five consecutive frames were included in the analysis and an R^2^ goodness-of-fit threshold above 0.8 was applied to ensure reliable growth rate estimations. The final dataset included growth rates, track IDs, and replicate identifiers.

### Neighbourhood analysis

To assess how the presence of neighboring mutants influences cell growth, we analyzed the spatial composition of each cell’s local environment. For every focal cell, we quantified the fraction of surrounding cells that were mutants based on pixel area. Specifically, we defined the mutant fraction as the area occupied by mutant cells divided by the total area of all cells (mutant + wildtype) within a circular neighborhood centered on the cell centroid, excluding the focal cell itself. Cell segmentation data from time-lapse microscopy of methionine, proline, and tryptophan auxotrophs and their wildtype counterparts were used for this analysis. To determine the appropriate spatial scale for this analysis, we performed a systematic evaluation across a range of neighborhood radii and computed the Spearman rank correlation coefficient (ρ) between the growth rate of mutant cells and the mutant fraction in their surroundings. This analysis revealed a negative correlation - indicating that higher local mutant density is associated with reduced growth - which varied in strength with neighborhood size. Since the optimal neighborhood size (i.e., radius with strongest negative correlation) differed across conditions and replicates, we standardized the analysis using a neighborhood radius of 5 µm, which represented the minimal interaction range consistently observed across all three amino acid auxotroph types. Final growth-versus-neighborhood analyses were performed using this 5 µm radius, and Spearman’s rank correlation was used to quantify the relationship between local mutant fraction and growth rate for both mutant and wildtype cells across time points.

### Statistical Analysis

All statistical analyses were conducted using Python version 3.11.4, utilizing built-in statistical functions from the scipy.stats package. To compare growth rates between wildtype and auxotrophic cells, we performed paired t-tests, using the arithmetic mean growth rate of each of the five independent replicates as individual data points. Kruskal-Wallis tests were used to evaluate overall differences in growth rate distributions between conditions. When external amino acids were provided, paired t-tests were used to test whether auxotrophs exhibited significantly higher growth rates compared to wildtypes. For all statistical tests, p-values were considered significant if they were below 0.05. Statistical significance is denoted as follows: *p* < 0.05 (^*^*), p < 0*.*01 (*^*\*\**^*), and p < 0*.*001 (*^*\*\*\**^).

To assess the relationships between single-cell growth rates and neighborhood composition, we computed Spearman’s rank correlation coefficients (ρ). This non-parametric test measures monotonic relationships without assuming linearity. The statistical significance of the observed correlations was determined through permutation testing, wherein the associations between growth rates and neighborhood compositions were randomized 10,000 times to generate a null distribution. The p-value was computed as the fraction of permutations that yielded correlation coefficients equal to or more extreme than the observed values.

To evaluate the effects of local mutant clustering on auxotroph growth, we computed the fraction of auxotrophic cells within 5, 9 and 13 µm neighborhood surrounding each focal cell and assessed its correlation with individual cell growth rates. Spearman correlation coefficients were calculated separately for each replicate, and the significance of the observed trends was assessed using permutation testing. For wildtype cells, a similar approach was used to determine whether the presence of neighboring auxotrophs impacted wildtype growth rates.

### Mutant specific growth rate prediction model

We adapted the mathematical model derived by van Vliet et al. (2022)^31^ to predict the relative growth rate of auxotrophic mutants compared to wildtype cells in dense, spatially structured populations. Specifically, this model provides an analytical expression for the highest possible growth rate an auxotroph can achieve when surrounded entirely by producing cells. Under these conditions, the auxotroph’s consumption of the essential amino acid has a negligible impact on its local concentration, allowing us to assume that the available amino acid is at equilibrium, as it would be in a space fully occupied by producing cells.

We parameterized this equation for the specific case of an auxotrophic mutant growing in a wildtype-dominated environment, where all surrounding cells are capable of producing the limiting amino acid. The highest possible growth rate (*r*) that the auxotroph can achieve, relative to the wildtype growth rate, is given by:

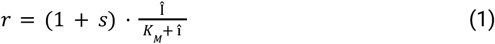

The first term represents the growth benefit the auxotroph obtains from not having to produce the essential amino acid. Specifically, the parameter *s* represents its fitness benefit and is defined as:

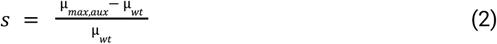

where μ_max,aux_ is the maximum growth rate of the auxotroph when the essential amino acid is externally supplied at a non-limiting concentration and μ_wt_ is the wildtype growth rate in minimal media, where wildtype cells must synthesize the essential amino acid.

The second term represents a Monod growth term assuming growth is limited by the internal concentration of the essential amino acid Î. It was previously shown^31^ that this is given by

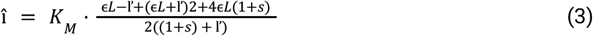

In this equation, *K*_*M*_ is the half-saturation (Monod) constant for growth on the limiting amino acid. The parameter ϵ represent the ratio of uptake (*u*_*aux*_, *u*_*wt*_) and leakage (*l*_*aux*_, *l*_*wt*_) rates of this amino acid between auxotrophs (aux) and wildtype (wt) cells, respectively:

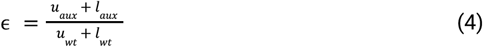

In the absence of gene regulatory changes in amino acid transporters or additional mutations affecting export/import pathways, we expect auxotrophs and wildtypes to have identical uptake and leakage rates, i.e. ϵ = 1.

The term ľ represents the relative leakage rate of auxotrophs relative to their maximum growth rate when amino acids are not growth limiting:

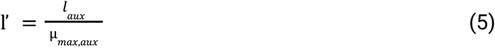

Finally, the parameter *L* represent the relative leakage flux which quantifies how much of the essential amino acid is externalized by wildtype cells compared to how much they use for biomass production:

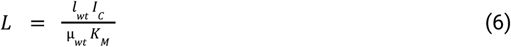

In this equation *I*_*C*_ is the internal concentration of the essential amino acid in the producing wildtype cells. We assume that wildtype cells can maintain a sufficiently high internal concentration to grow close to their maximum rate, and thus assume that *I*_*C*_ = 20 · *K*_*M*_.^10^

### Inference of leakage rates

To estimate leakage rates from the imaging data, we fitted Eq. 1 to the experimentally determined maximum growth rate of auxotrophs in wildtype-dominated environments. Specifically, we extrapolated the linear regression of auxotroph growth rate as a function of the fraction of wildtype cells in the local neighborhood to a wildtype frequency of 1. This extrapolated value corresponds to the maximum growth rate that an auxotroph can achieve when fully surrounded by wildtype cells. We then fitted the model by setting l_aux=l_wt=L such that both cell types share the same leakage parameter. Because Eq. 1 depends monotonically on the leakage rate parameter L, we determined the leakage rate by numerically identifying the value of L at which the predicted maximum growth rate matched the experimentally estimated maximum. This approach provides an order-of-magnitude estimate of leakage rates in structured populations, while acknowledging that precise biochemical quantification would require dedicated assays beyond the scope of this study.

Although the inferred leakage rates are sensitive to model assumptions, our aim was not to obtain absolute values but to position our experimental system within the model’s parameter space. Because both inference and predictions rely on the same assumptions, systematic biases tend to cancel out, enabling meaningful comparisons between strains even if absolute parameter values shift.

## Limitations of the Study

The design of and conditions in our microfluidic chambers restrict where stable clusters can form. Cells in the chamber center are continuously displaced and washed out, so only cells located at the sidewalls persist and can form clusters. Consequently, clustering effects are measured mainly at chamber boundaries and may underestimate dynamics in environments where cells can accumulate throughout a habitat.

Our experiments also focus on short-term interactions within a single *E. coli* species under balanced exponential growth. They do not incorporate environmental fluctuations, nutrient limitation, or cell lysis, all of which can trigger active metabolite release and alter amino acid availability. Moreover, natural microbial communities contain multiple species with distinct leakage rates, uptake efficiencies, and metabolic strategies. Such multispecies effects which can create or eliminate ecological niches for auxotrophs will be interesting to investigate in future studies.

Finally, our mathematical model simplifies spatial metabolite diffusion and relies on average leakage and uptake parameters. While this abstraction captures the key trade-offs, it does not represent local heterogeneity in cell placement, diffusion barriers, or stochastic fluctuations in metabolite pools that may influence auxotroph fitness in more complex environments.

## Supplemental Information

Document S1. Figures S1 to S8, Table S1 and S2, and supplemental references.

## Resource availability

### Lead contact

Requests for further information and resources should be directed to and will be fulfilled by the lead contact, Martin Ackermann.

### Materials availability

This study did not generate new unique reagents, plasmids or strains.

### Data availability

- All the raw imaging data generated in this study have been deposited in the Eawag Research Data Institutional Collection (ERIC) and are publicly available as of the date of publication at https://doi.org/10.25678/000FSR.
- All original code has also been deposited in the Eawag Research Data Institutional Collection (ERIC) and are publicly available as of the date of publication at https://doi.org/10.25678/000FSR.
- All other items - Any other information required to reanalyze the data reported in the paper is available from the lead contact upon request.

## Key Resources Table

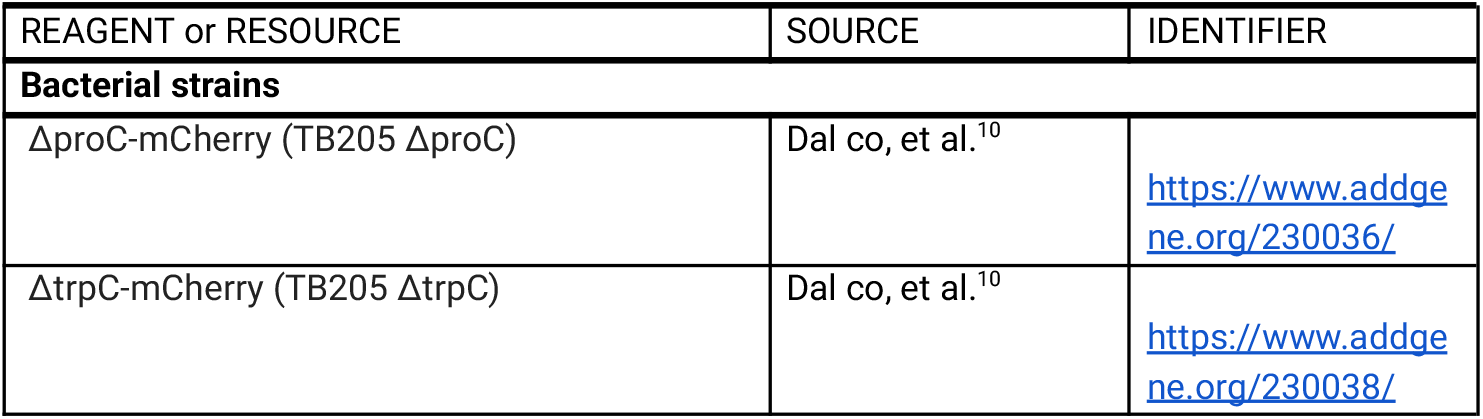
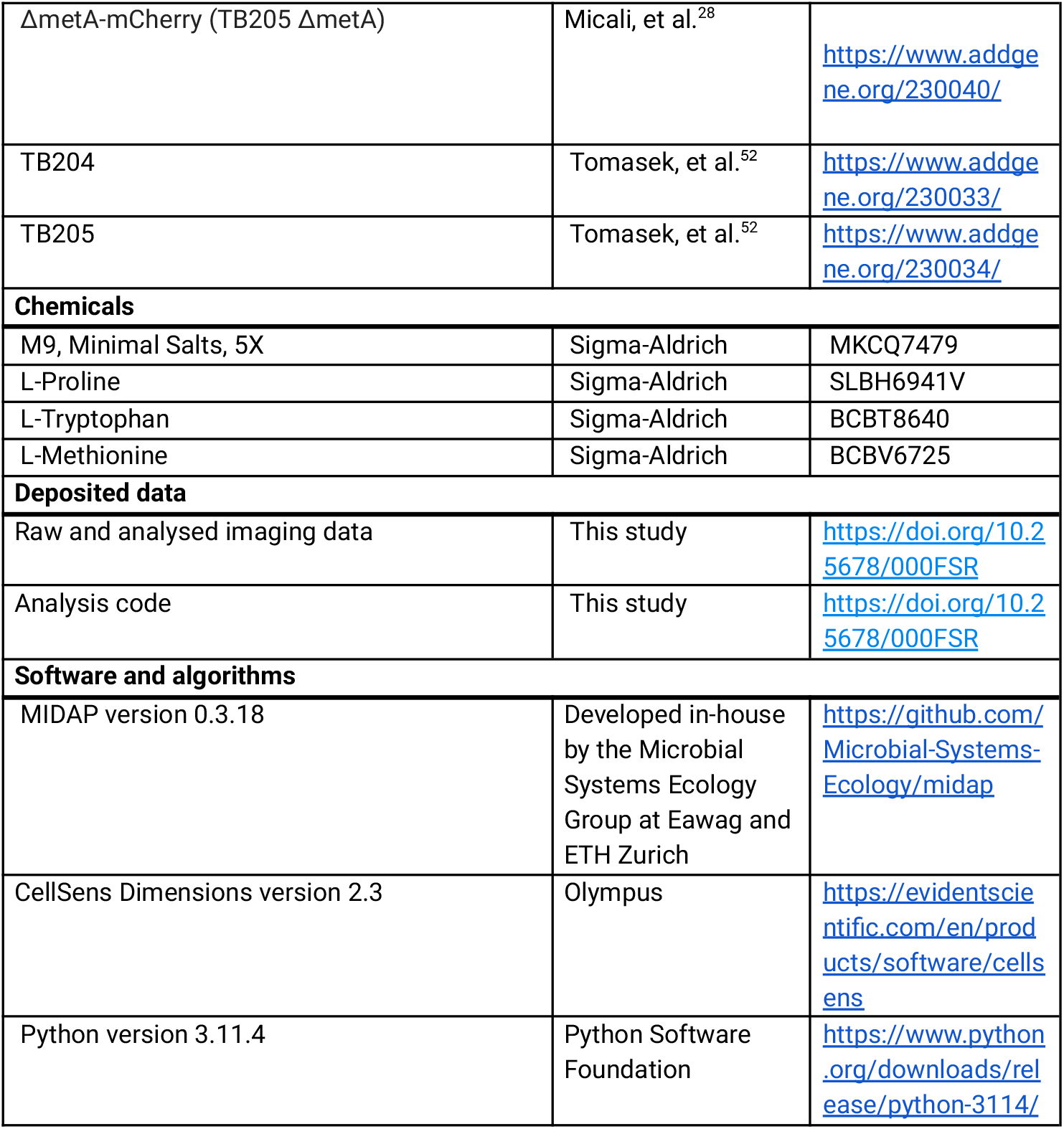

| REAGENT or RESOURCE | SOURCE | IDENTIFIER |
| --- | --- | --- |
| **Bacterial strains** |  |  |
| ΔproC-mCherry (TB205 ΔproC) | Dal co, et al.<sup>10</sup> | https://www.addgene.org/230036/ |
| ΔtrpC-mCherry (TB205 ΔtrpC) | Dal co, et al.<sup>10</sup> | https://www.addgene.org/230038/ |
| ΔmetA-mCherry (TB205 ΔmetA) | Micali, et al.<sup>28</sup> | https://www.addgene.org/230040/ |
| TB204 | Tomasek, et al.<sup>52</sup> | https://www.addgene.org/230033/ |
| TB205 | Tomasek, et al.<sup>52</sup> | https://www.addgene.org/230034/ |
| **Chemicals** |  |  |
| M9, Minimal Salts, 5X | Sigma-Aldrich | MKCQ7479 |
| L-Proline | Sigma-Aldrich | SLBH6941V |
| L-Tryptophan | Sigma-Aldrich | BCBT8640 |
| L-Methionine | Sigma-Aldrich | BCBV6725 |
| **Deposited data** |  |  |
| Raw and analysed imaging data | This study | https://doi.org/10.25678/000FSR |
| Analysis code | This study | https://doi.org/10.25678/000FSR |
| **Software and algorithms** |  |  |
| MIDAP version 0.3.18 | Developed in-house by the Microbial Systems Ecology Group at Eawag and ETH Zurich | https://github.com/Microbial-Systems-Ecology/midap |
| CellSens Dimensions version 2.3 | Olympus | https://evidentscientific.com/en/products/software/cellsens |
| Python version 3.11.4 | Python Software Foundation | https://www.python.org/downloads/release/python-3114/ |

## Acknowledgements

We thank Snorre Sulheim and Gabriele Micali for valuable discussions on experiments and data analysis. We are also grateful to the members of the Microbial Systems Ecology group for insightful discussions throughout this work. This research was supported by ETH Zurich and Eawag, the Swiss Federal Institute of Aquatic Science and Technology. Additionally, it was funded as part of the NCCR Microbiomes, supported by the Swiss National Science Foundation (grant numbers 51NF40_180575 and 51NF40_225148), and through a project grant from the Swiss National Science Foundation (188642). SvV was additionally supported by an Ambizione grant (grant no. 202186) from the Swiss National Science Foundation and by the University of Basel.

## Author contributions

Conceptualization and design of the study: DR, OS, MA

Experimental data collection: DR, EF, SU

Data analysis: DR, FO, SvV

Funding acquisition: MA, OS

Project management: DR

Supervision: MA, OS, SvV

Writing – original draft: DR

Writing – review & editing: DR, OS, MA and all other co-authors

## Declaration of the use of generative AI and AI-assisted technologies in the manuscript preparation process

During the preparation of this work, the authors used ChatGPT (versions 4o and 5) to assist with text rephrasing and code debugging for data analysis prior to submission. All outputs generated by the tool were subsequently reviewed and edited by the authors, who take full responsibility for the final content of the publication.

