## Supplementary Information for "Limits to the evolution of metabolic dependency in spatially structured microbial communities"

### Supplementary Figures

**Figure S1. Replicate-wise growth rate distributions and statistical comparisons of wildtype and auxotrophic mutant cells in the absence and presence of amino acids.** Violin plots displaying growth rate distributions for wildtype and mutant cells across individual replicates for methionine, proline, and tryptophan auxotroph-wildtype experiments in the absence (a) and presence (b) of amino acids . This figure provides the replicate-level breakdown underlying the data shown in *Figure 1*.Kruskal-Wallis tests were performed to assess whether there are statistically significant differences between wildtype and auxotroph growth rates across all replicates. In the absence of amino acids, growth rate differences were significant for methionine (H = 2451.75, $p < 0.001$ ), proline (H = 2502.51, $p < 0.001$), and tryptophan (H = 2490.84, $p < 0.001$). Similarly, in the presence of amino acids, significant differences were observed for methionine (H = 67.67, $p < 0.001$), proline (H = 265.89 $p < 0.001$), and tryptophan (H = 109.25, $p < 0.001$).

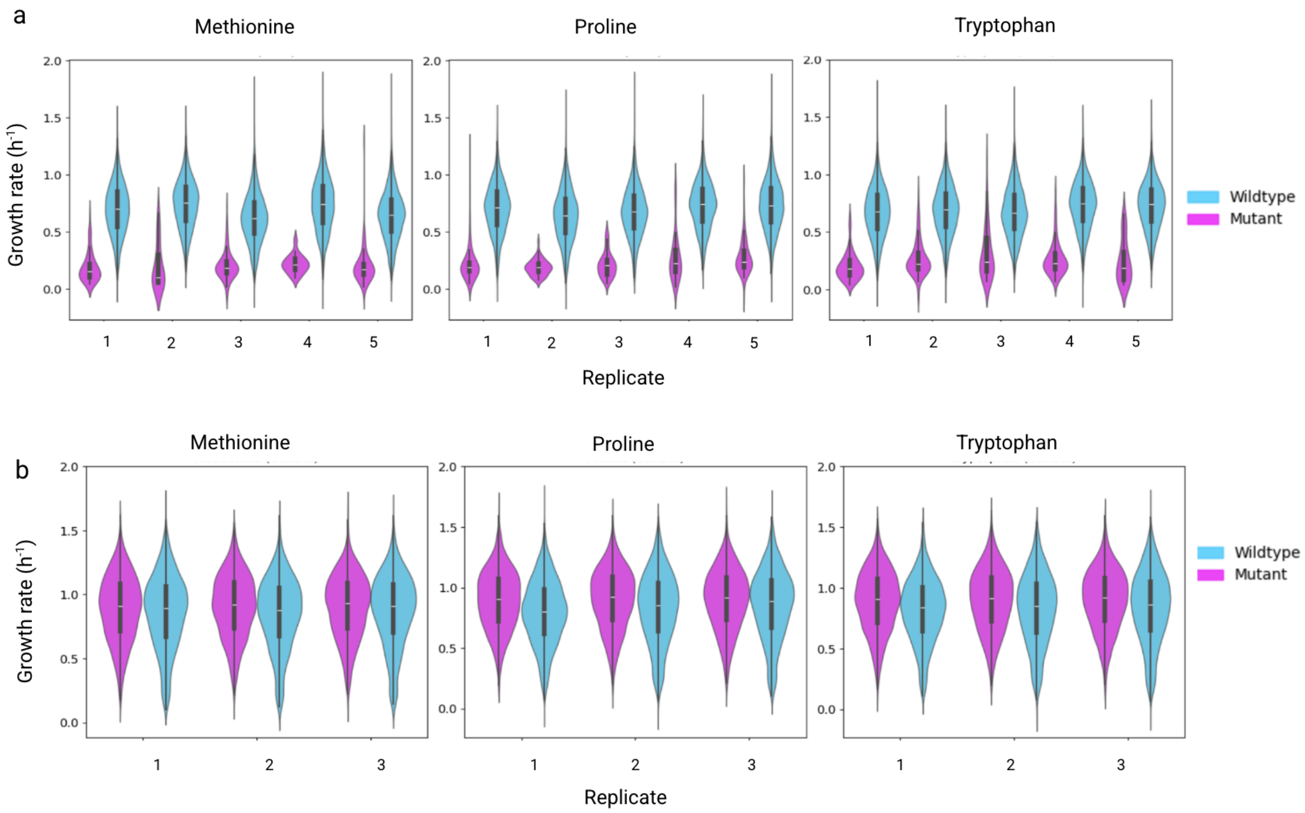

**Figure S2. Relationship between local neighborhood composition and growth rates across replicates in the absence of amino acids.** Scatter plots illustrating how the fraction of mutants in a 5-micrometer radius around a focal cell affects its growth rate in the absence of amino acids for mutant cells (a) and wildtype cells (b). Each data point represents a single cell. This figure provides the replicate-level breakdown underlying the data shown in *Figure 2*. Box plots overlaid on the scatter illustrate the distribution of growth rates across fixed bins of the local auxotroph fraction. While binning was applied for visualization, all statistical analyses were performed on the original, continuous data.

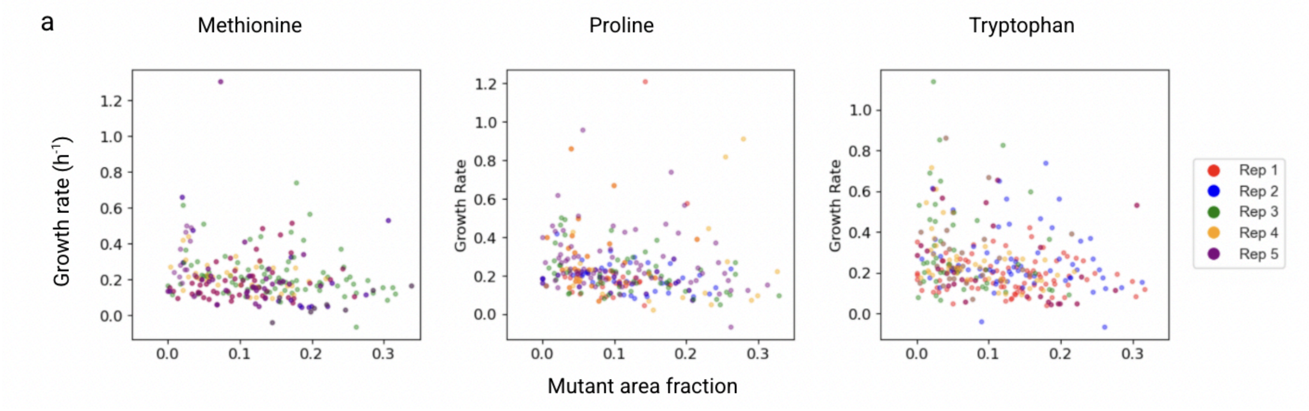


Spearman correlation coefficients for each of the five independent replicate experiments:

- Methionine: -0.41, -0.18, -0.35, -0.10, -0.30
- Proline: -0.28, -0.15, -0.40, -0.19, -0.32
- Tryptophan: -0.48, -0.16, -0.37, -0.24, -0.27

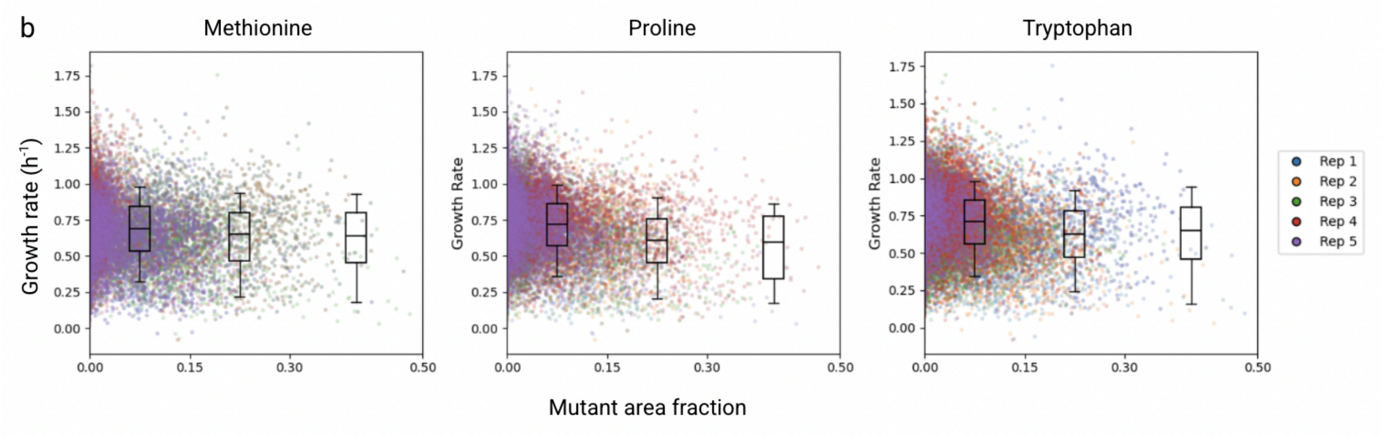


Spearman correlation coefficients for each of the five independent replicate experiments:

- Methionine: -0.411, -0.184, -0.352, -0.095, -0.297
- Proline: -0.275, -0.146, -0.401, -0.194, -0.318
- Tryptophan: -0.475, -0.164, -0.368, -0.240, -0.271

**Figure S3. Permutation tests for Spearman correlations between local mutant fraction and growth rates in the absence of amino acids.** Permutation tests were performed to assess the statistical significance of the Spearman correlations between local mutant fraction and growth rate of a focal cell in the absence of amino acids as shown in *Figure S2*. To generate a null distribution, we randomized the data by scrambling the associations between local mutant fraction and growth rate while preserving the overall distribution of each variable. Spearman correlation coefficients from the real data (vertical line, each representing a biological replicate) were then compared to this null distribution to evaluate significance. (a) For mutants, the observed negative correlations between local mutant fraction and growth rate fall outside the distribution generated from randomized data, indicating statistical significance (typically $p < 0.001$, with one methionine replicate significant at $p < 0.05$). This suggests that higher mutant fractions in the neighborhood are associated with slower mutant growth. (b) Similarly, for wildtype cells, we observed statistically significant negative correlations ($p < 0.05$) between local mutant fraction and growth rate. These findings suggest that mutant presence exerts a suppressive effect not just on clonal mutant populations, but also on neighboring wildtype cells.

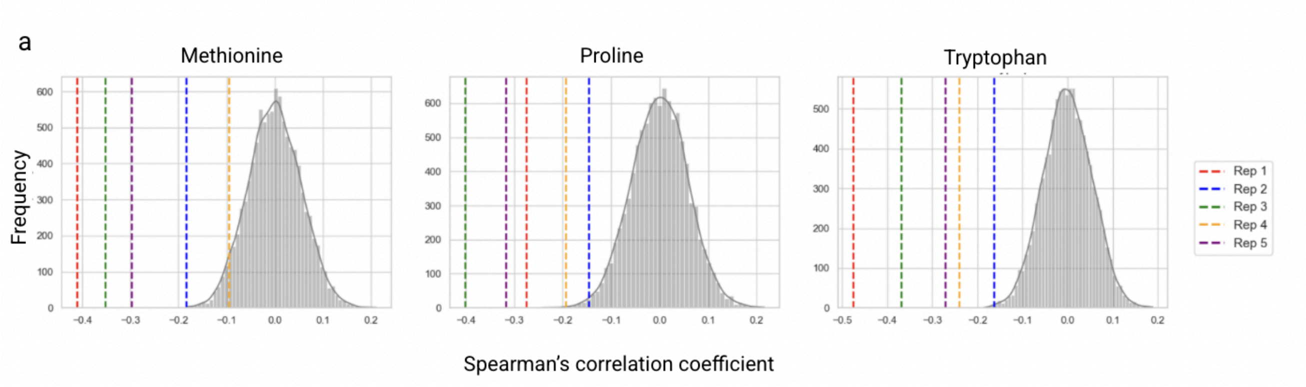


P-values for each of the five independent replicate experiments were less than 0.05.

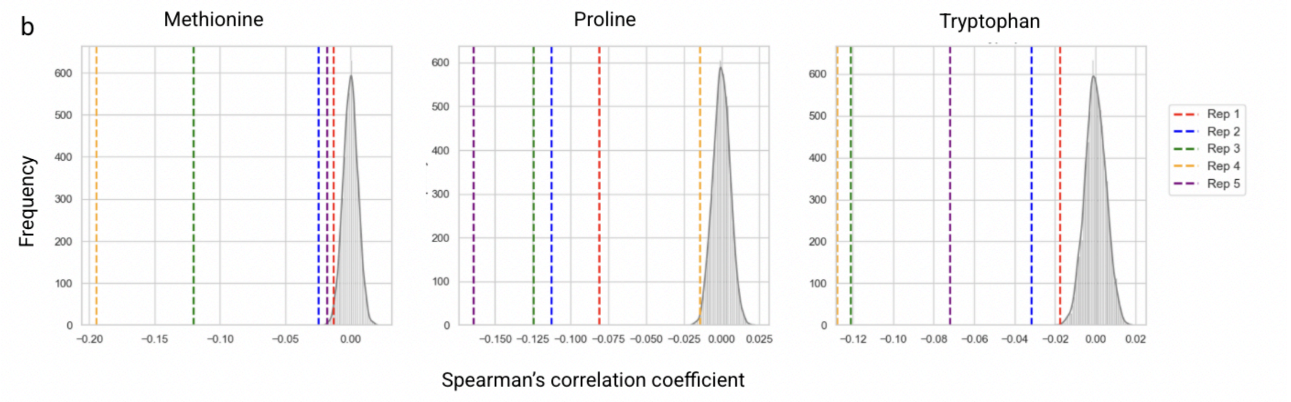


P-values for each of the five independent replicate experiments were less than 0.05.

**Figure S4. Sensitivity analysis of the relationship between mutant growth rate and mutant area fraction within 5 µm, 9 µm and 13 µm radii in the absence of amino acids.** Scatter plots of mutant growth rate versus mutant area fraction for tryptophan, methionine and proline auxotrophs at three neighborhood radii: 5 µm, 9 µm and 13 µm. Each data point represents a single cell. Box plots show the distribution of growth rates across fixed bins of the local auxotroph fraction. While the data was binned for visualization purposes, all statistical tests (Spearman correlation and permutation tests, not shown) were performed on the original continuous data. Auxotrophs surrounded by a higher fraction of other auxotrophs exhibit significantly reduced growth, reflecting negative selection that intensifies with local auxotroph frequency. In all cases the Spearman rank-correlation between mutant growth rate and mutant area fraction was significant ($p$ < 0.05; not shown).

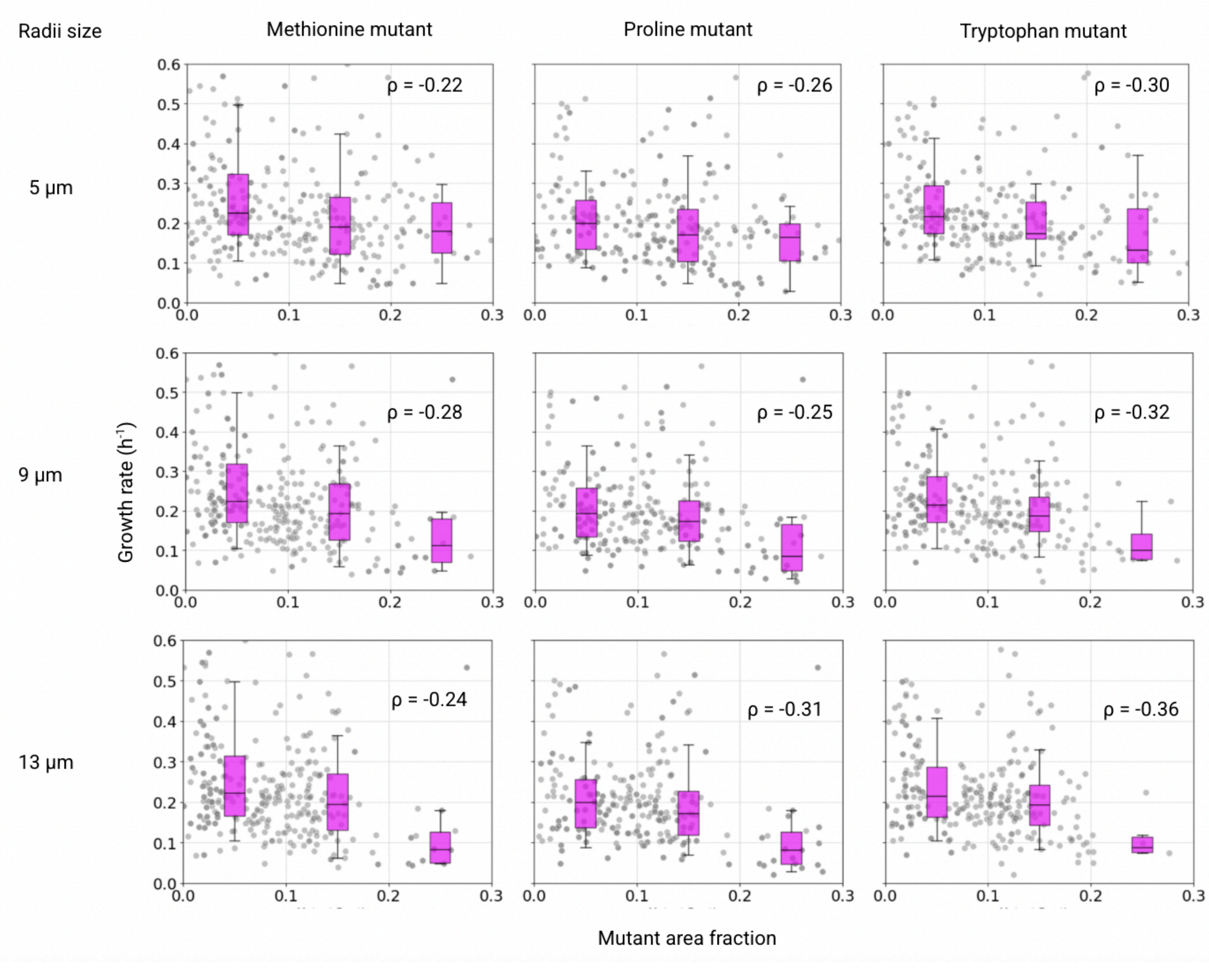

**Figure S5. Relationship between mutant area fraction and mutant growth rates across replicates in the presence of amino acids.** Scatter plots illustrating how the local neighborhood composition affects the growth rates of both mutant and wildtype cells in the presence of amino acids. (a) Mutant growth rate vs. local mutant area fraction, where each point represents a single auxotrophic mutant cell. (b) Wildtype growth rate vs. local mutant area fraction, where each point represents a single wildtype cell.

a

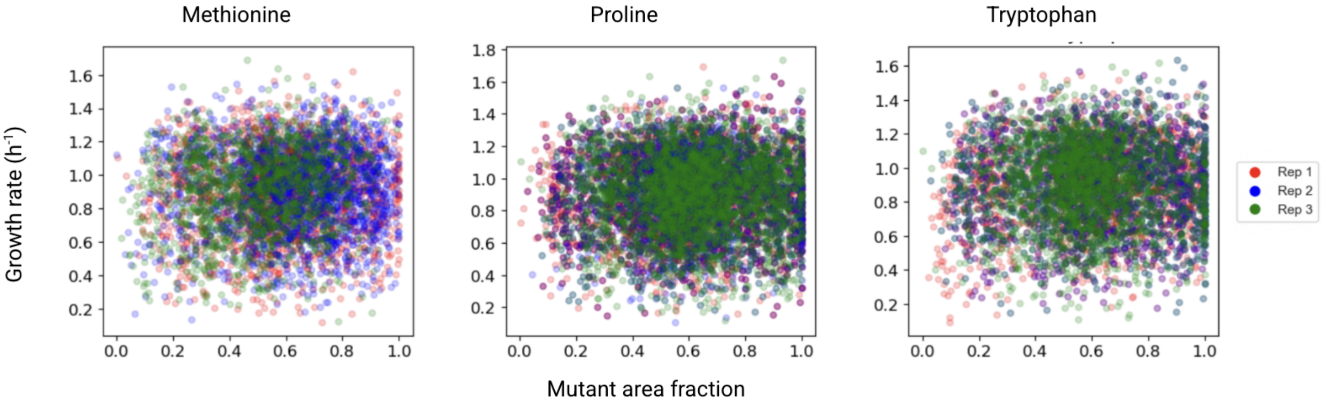


Spearman correlation coefficients for each of the three independent replicate experiments:

- Methionine: 0.026, 0.127, 0.024
- Proline: 0.007, 0.001, 0.015
- Tryptophan: -0.003, 0.070, -0.010

b

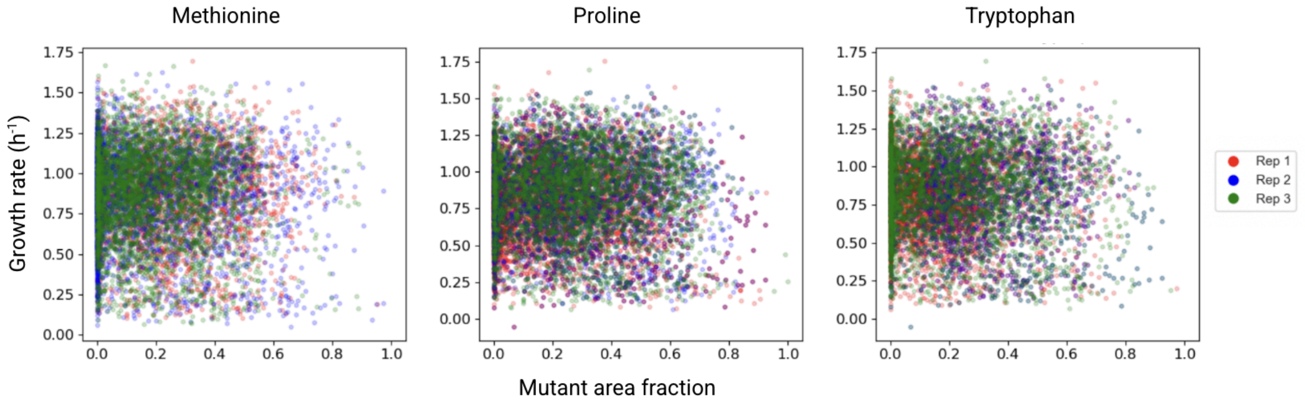


Spearman correlation coefficients for each of the three independent replicate experiments:

- Methionine:0.026, 0.008, 0.029
- Proline: 0.010, -0.000, 0.010
- Tryptophan: -0.006, 0.001, -0.011

**Figure S6. Permutation tests for Spearman correlations between local mutant fraction and growth rates in the presence of amino acids.** Permutation tests were performed to assess the statistical significance of the Spearman correlations between local mutant fraction and the growth rate of a focal cell in the presence of amino acids, as shown in *Figure S5*. To generate a null distribution, we randomized the data by scrambling the associations between local mutant fraction and growth rate while preserving the overall distribution of each variable. Spearman correlation coefficients from the real data (vertical lines, each representing a biological replicate) were then compared to this null distribution to evaluate significance. (a) For mutants, the observed correlations between local mutant fraction and growth rate lie within the distribution generated from randomized data ($p > 0.05$ for all replicates), indicating no significant relationship. This suggests that in the presence of externally supplied amino acids, mutant growth is unaffected by local neighborhood composition. (b) Similarly, for wildtypes, the observed correlation coefficients fall within the distribution from randomized data ($p > 0.05$ for all replicates), suggesting that wildtype growth is also independent of the fraction of mutants in the surroundings and is primarily driven by the external amino acid supply.

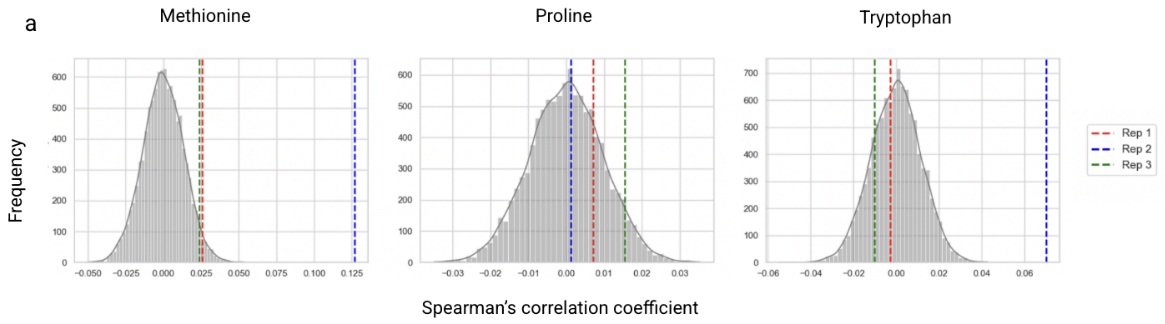


P-values for each of the three independent replicate experiments:

- Methionine: 0.464, 0.299, 0.477
- Proline: 0.278, 0.350, 0.232
- Tryptophan: 0.438, 0.054, 0.374

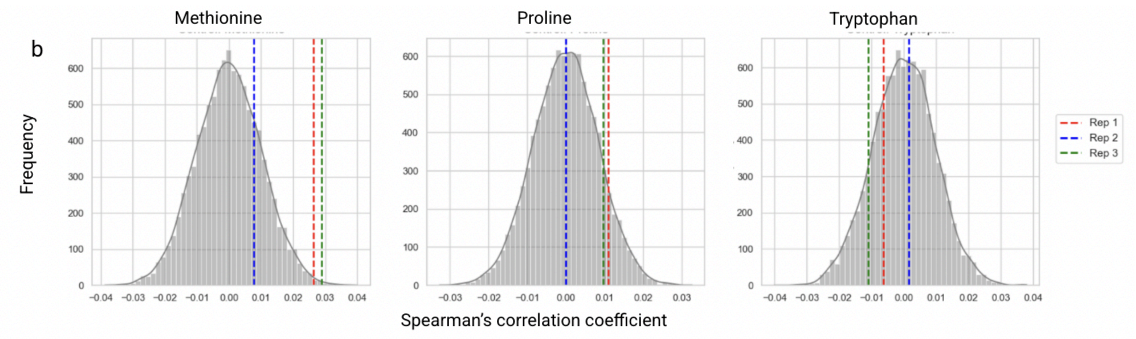


P-values for each of the three independent replicate experiments:

- Methionine: 0.995, 0.768, 0.998
- Proline: 0.875, 0.495, 0.874
- Tryptophan: 0.563, 0.259, 0.130

**Figure S7. Auxotrophic mutants are unable to invade wildtype populations.** Cell counts of auxotrophic mutants in co-cultures in the absence of externally supplied amino acids were determined over time. The frequency of mutants decreases over time, indicating that the selective disadvantage of auxotrophic mutants leads to an inability to invade the wildtype population. Lines show individual replicates, and shaded regions represent the confidence interval.

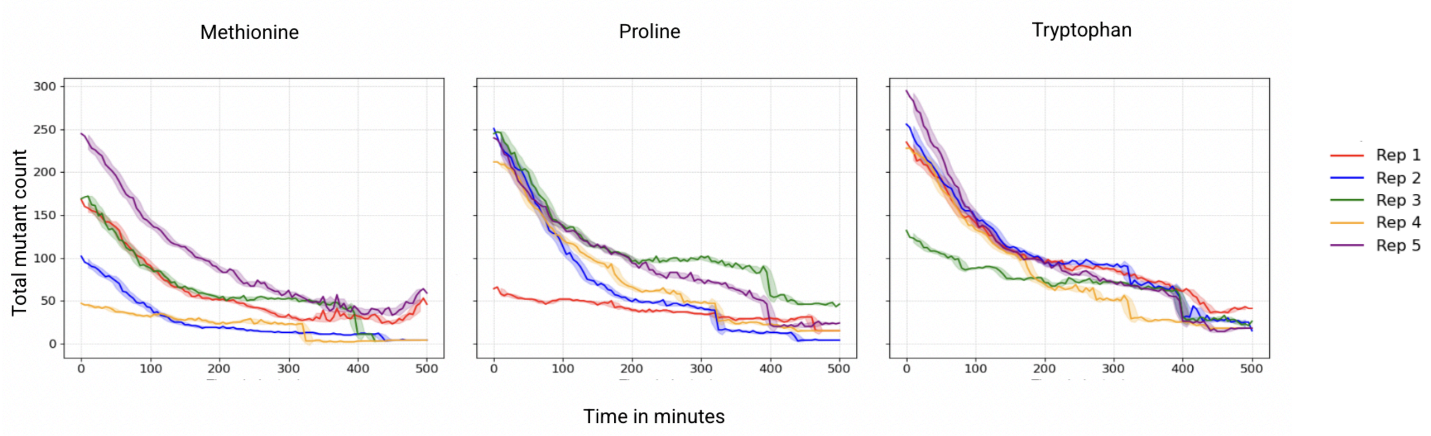


**Figure S8. GFP and mCherry marker gene expression does not affect growth differentially.** Growth of the TB204 and TB205 strains expressing GFP and mCherry marker genes, respectively, was measured in M9 minimal medium with amino acids (a) and without amino acids (b) using a plate reader (n = 3 biological replicates per strain). Curves show mean ± SD of $OD_{600}$ over time. In the presence of amino acids, the average maximal growth rates were 0.338 ± 0.01 $h^{-1}$ for TB204 and 0.351 ± 0.02 $h^{-1}$ for TB205, corresponding to a 3.8% difference (t = 0.541, $p$ = 0.612). In the absence of amino acids, the average maximal growth rates were 0.350 ± 0.01 $h^{-1}$ for TB204 and 0.345 ± 0.02 $h^{-1}$ for TB205, corresponding to a 1.3% difference (t = 0.500, $p$ = 0.643). Under both conditions, growth rates were nearly identical, indicating that the fluorescent marker genes do not differentially affect growth.

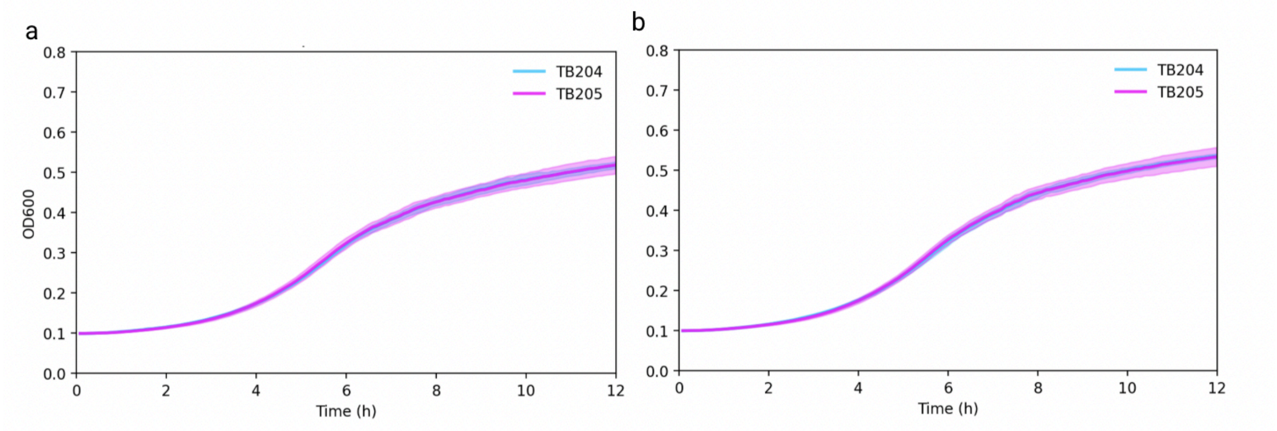

### Supplementary Tables

**Table S1. Model parameters summary.** This table summarises the key model parameters, including amino acid leakage rates[1], uptake rates[2,3], internal concentrations[4], growth benefits, and diffusion constants[5] for all the auxotrophs. Growth benefits of auxotrophy and wildtype growth rate were experimentally quantified in this study, while the leakage rates were fitted to the data; all other parameters were obtained from the literature.

| Parameter | Symbol used in the equations | Methionine (ΔmetA) | Proline (ΔproC) | Tryptophan (ΔtrpC) |
|---|---|---|---|---|
| Leakage rate [1/s] | l | $6.5 \times 10^{-7}$ [$4.7 \times 10^{-7}$ – $8.8 \times 10^{-7}$ 95% CI] | $1.5 \times 10^{-6}$ [$1.2 \times 10^{-6}$ – $1.9 \times 10^{-6}$ 95% CI] | $9.7 \times 10^{-7}$ [$8.6 \times 10^{-7}$ – $1.4 \times 10^{-6}$ 95% CI] |
| Uptake rate [1/s] | u | 6.30 | 2.03 | 24.04 |
| Growth benefit of auxotrophy [fold-change in specific growth rate] | s | 1.3 ± 0.02 | 1.26 ± 0.01 | 1.27 ± 0.01 |
| Wildtype growth rate [1/h] | $\mu_{wt}$ | 0.9 | 0.9 | 0.9 |
| Diffusion constant [$\mu m^2/s$] | D | 785 | 818 | 671 |

**Table S2. Summary statistics of single-cell growth rates.** For each condition, amino acid, and cell type, we report the number of replicates and the number of cells analyzed overall, along with their mean specific growth rate, standard deviation, standard error of the mean.

Without amino acids:

| Amino acid | Cell type | Number of cells analysed | Mean specific growth rate | Standard deviation | Standard error of the mean | Number of biological replicates |
|---|---|---|---|---|---|---|
| Methionine | mutant | 337 | 0.20 | 0.14 | 0.008 | 5 |
| Methionine | wildtype | 30879 | 0.68 | 0.22 | 0.001 | 5 |
| Proline | mutant | 295 | 0.24 | 0.15 | 0.008 | 5 |

| | | | | | | |
|---|---|---|---|---|---|---|
| Proline | wildtype | 30444 | 0.71 | 0.22 | 0.001 | 5 |
| Tryptophan | mutant | 346 | 0.25 | 0.16 | 0.008 | 5 |
| Tryptophan | wildtype | 36748 | 0.70 | 0.21 | 0.001 | 5 |

With amino acids:

| **Amino acid** | **Cell type** | **Number of cells analysed** | **Mean specific growth rate** | **Standard deviation** | **Standard error of the mean** | **Number of biological replicates** |
|---|---|---|---|---|---|---|
| Methionine | mutant | 5513 | 0.90 | 0.25 | 0.003 | 3 |
| Methionine | wildtype | 9775 | 0.86 | 0.28 | 0.002 | 3 |
| Proline | mutant | 10828 | 0.90 | 0.25 | 0.002 | 3 |
| Proline | wildtype | 13894 | 0.82 | 0.28 | 0.002 | 3 |
| Tryptophan | mutant | 7482 | 0.90 | 0.25 | 0.002 | 3 |
| Tryptophan | wildtype | 10605 | 0.82 | 0.28 | 0.002 | 3 |
